# Simulations of fluorescence imaging in the oral cavity

**DOI:** 10.1101/2021.03.31.437770

**Authors:** Zheng Lyu, Haomiao Jiang, Feng Xiao, Jian Rong, Tingcheng Zhang, Brian Wandell, Joyce Farrell

## Abstract

We describe an end-to-end image systems simulation that models a device capable of measuring fluorescence in the oral cavity. Our software includes a 3D model of the oral cavity and excitation-emission matrices of endogenous fluorophores that predict the spectral radiance of oral mucosal tissue. The predicted radiance is transformed by a model of the optics and image sensor to generate expected sensor image values. We compare simulated and real camera data from tongues in healthy individuals and show that the camera sensor chromaticity values can be used to quantify the fluorescence from porphyrins relative to the bulk fluorescence from multiple fluorophores (elastin, NADH, FAD, and collagen). Validation of the simulations supports the use of soft-prototyping in guiding system design for fluorescence imaging.

© 2021 Optical Society of America under the terms of the OSA Open Access Publishing Agreement

## 1. Introduction

Implementing an imaging system for a domain-specific application requires selecting and integrating many different hardware and software components. An end-to-end simulation that models each component and how they work together can accelerate innovation by shortening the time-consuming and expensive design-build-test loop. We are developing and validating image systems simulation tools for soft-prototyping imaging systems for several domains, including consumer photography [1–3], underwater imaging [4], AR/VR displays [5] and autonomous driving [6–10].

In this paper we implement simulations to prototype a system for imaging and quantifying fluorescence in the oral cavity. The motivation for designing the system is based on observations that tissue autofluorescence can discriminate between normal and precancerous tissue [11–13]. This finding has led to the development of several different types of imaging systems designed for non-invasive in-vivo measurements of tissue autofluorescence [14]. Because the autofluorescence signal is weak compared to reflected light, one must select special purpose components that can separate reflected and fluorescent photons.

The image systems simulations enable us to evaluate different combinations of illuminants, filters and sensors that excite biological tissue fluorophores and quantify the photons emitted by the fluorophores. We validate simulations by comparing the prototype predictions with measurements from a real system that is designed to evoke fluorescence from the oral cavity. The soft-prototyping tools are sufficiently accurate to form the foundation for future work that explores alternative image system designs.

## 2. Background

Several types of fluorescent emissions in the oral cavity have been measured during precancerous and cancerous stages. These emissions have been measured using fiber optic illumination and spot spectroradiometric sensors [12, 15, 16]. The measurements reveal a complex set of changes in the oral cavity fluorophores in the presence of precancerous and cancerous tissue.

- Certain types of oral cavity tissue fluorescence are reduced in the presence of precancerous and cancerous tissue. The reduced fluorescence have been attributed to a reduction in FAD (flavin adenine dinucleotide), a molecule that plays an important role in cell respiration and metabolism, and to changes in collagen and elastin that occur with cellular damage [17–21].
- Other types of tissue fluorescence in the oral cavity are higher than normal in the presence of cancerous tissue. Several investigators have hypothesized that NADH fluorescence (the reduced form of nicotinamide adenine dinucleotide) increases in cancerous tissue [19, 22].
- Investigators also report observing the distinctive spectral signature of porphyrin fluorescence in cancerous lesions [12, 17, 23–26]. Porphyrin fluorescence is present in the mouths of many healthy individuals as well, and it can be measured in plaque [27], caries [28] and on the dorsal side of healthy tongues [20]. The porphyrin fluorescence is large, but not necessarily diagnostic of oral cancer [23].

The complex set of findings led to the development of several illumination systems designed to help dentists visualize oral mucosal abnormalities (e.g Velscope^®^, OralID^®^, Identafi^®^). These products use short-wavelength LEDs to excite endogenous fluorophores in the oral mucosal tissue. The clinician is provided with glasses or a viewer that block the reflected short-wavelength light from the illuminant and enhance the visibility of the fluorescent emissions in the middle- and long-wavelengths [13, 29, 30]. The clinician is tasked with judging whether there are abnormally dark areas on the tongue and in other parts of the oral cavity.

In practice, the size of the measured fluorescent signal depends markedly on the choice of illuminant. Many empirical reports use a single, narrowband illuminant to excite fluorescence with peak wavelengths ranging between 350 and 450 nm. One of the objectives of soft prototyping is to explore the consequences of selecting different combinations of illuminants and sensors.

Further, it is desirable to build an oral cancer screening system based on a quantitative lab test, rather than the clinician’s visual judgment. A system that reliably measures fluorescence may provide the basis of a lab test that meaningfully assesses the health status of the oral cavity. A second objective is to establish how well the image system can quantify fluorescence from the oral cavity.

## 3. Methods: Image Systems Simulation

The image systems soft-prototyping tools are based on a quantitative model of the scene and image acquisition device. Implementing the simulation requires defining: (a) a three-dimensional graphics model of the key elements (oral cavity, light and camera positions), (b) the lights and materials, including their spectral reflectance and fluorescence, and (c) a model of the camera, including its optics and sensor.

### 3.1. Geometry of the oral cavity, light, and camera

We use computer graphics packages, including Cinema 4D and Blender, to represent the geometry of the scene (Figure 1(a)). This application requires geometric modeling of the size and shape of the oral cavity, the positions of the imaging system’s lights, filters, lens and sensor. The oral cavity represents the surfaces (tongue, lips, teeth, etc.) as meshes and the relative positions of the image system components as points or regions in 3d-coordinates. The geometric data are exported to a set of text files that are read and interpreted by the physically-based ray-tracing software (PBRT, [31]). The physical properties of the lights, materials, and lens are specified using the parameters of the PBRT software (Figure 1(b)). Some of the critical properties, such as specifying tissue fluorescence or absolute spectral power distributions of the illumination, were enabled by open-source modifications that we added to the open-source PBRT code for this project. These features are included in the freely available Docker container used to create the simulations in this paper.

**Fig. 1.**
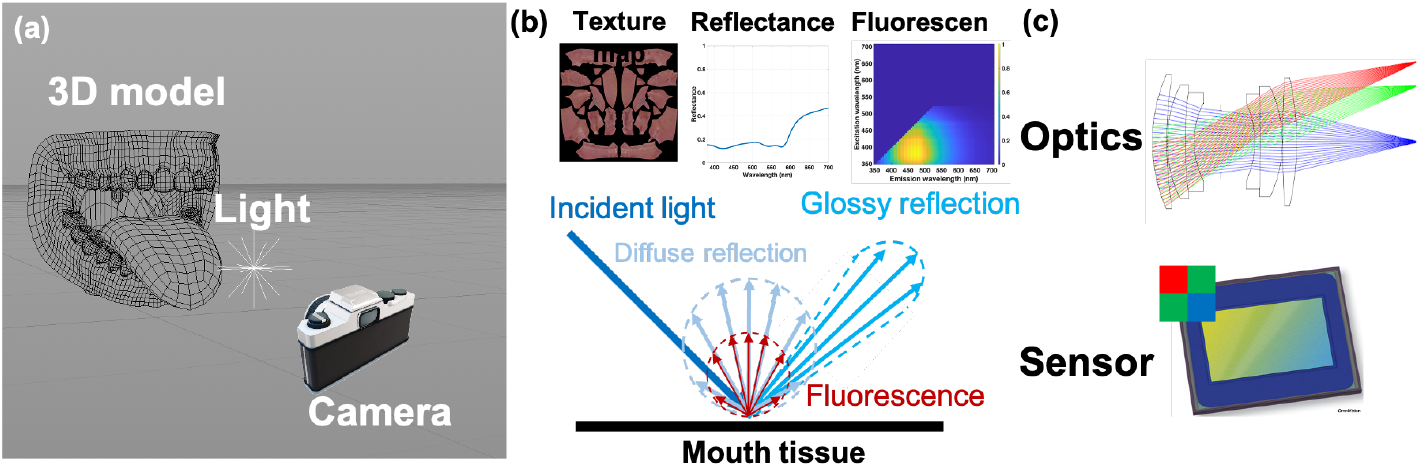
Diagram of the imaging system simulation pipeline. a) The 3D mesh model of the oral cavity, as well as the positions of the light and camera, are defined in graphics software. b) The simulations incorporate models of the tongue texture map, surface reflectance, and tissue fluorescence. The ray tracing also models diffuse and glossy reflections. c) The camera model specifies the multi-element optics as well as the spectral quantum efficiency, geometric and electrical properties of the image sensor.

Controlling the physical properties of the materials (e.g., fluorophore concentration, diffuse reflectance, spectral power distribution, light intensities) is an essential part of the simulation environment. We used the Matlab toolbox (ISET3d) to simplify programmatic control of the assets, materials, textures, and illumination^1^ [7]. The toolbox includes functions that read the PBRT text files, represent them as internal Matlab objects, and enable the user to set and get properties of the entire scene. The toolbox also includes functions that save out the modified parameters in PBRT format and then invoke the Docker container with the PBRT ray-tracer to render the scene spectral radiance or the sensor image irradiance. The ability to programmatically control the scene properties is essential as we test different systems and a range of measurement conditions including (a) different poses of the tongue within the oral cavity, and (b) different positions of the lights and camera with respect to the oral cavity. Figure 2 illustrates examples of different tongue and jaw poses that can be programmatically controlled as part of the simulation.

**Fig. 2.**
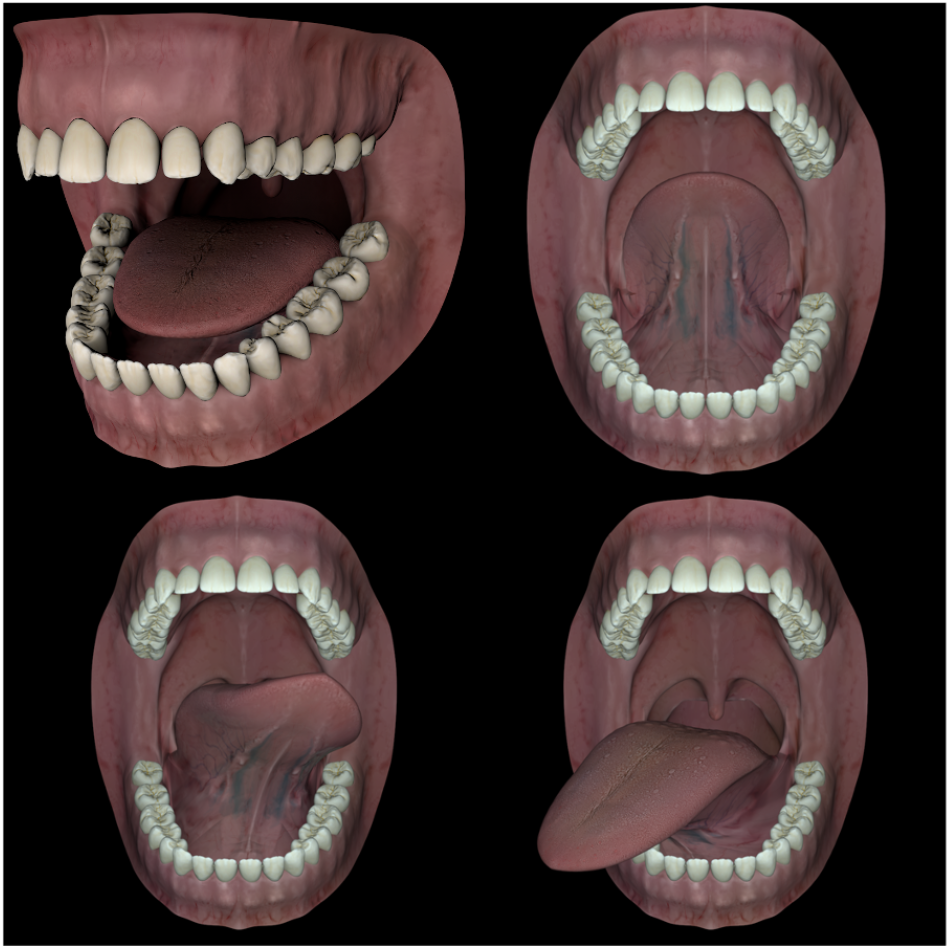
Mouth model rendered in various poses.

### 3.2. Material scattering and fluorescence

The spectral radiance *r*(λ) from the tissue surface can be partitioned into two additive components, the standard diffuse-glossy reflection *r*_*ref*_ and a fluorescent emission *r* _*fluo*_:

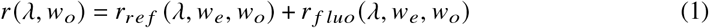

In the ray-tracing protocol of PBRT the diffuse-glossy reflectance and fluorescent emission of the scene radiance are parameterized by the angles of the incident ray (*w*_*e*_) and the outgoing ray (*w_o_*). The radiance from the diffuse-glossy reflectance is calculated as the wavelength-by-wavelength product of the irradiance, *L* (*λ,w*_*e*_), and the surface reflectance, *R*(*λ,w*_*e*_, *w*_*o*_):

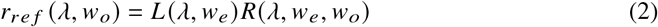

The calculations of these angle-dependent ray intensities rely on standard material models, in this case a combination of a Lambertian term and a small glossy term that gives the tongue and teeth their shiny appearance.

The fluorescence properties of the material is characterized by an excitation emission function *e*(*λ, λ*_*i*_), where *λ* is the incident light wavelength and is the fluorescence emission wavelength. The calculation of the fluorescence is the product of the spectral irradiance with the excitation-emission matrix. Specifically, we calculate the fluorescent emission at *λ*, given an incident ray at *λ*_*i*_ and angle *w*_*e*_, using:

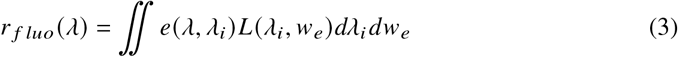

We model the angle of the fluorescent output as Lambertian, so that no term is needed (all angles are equal). The excitation and emission function for each fluorophore is also referred to as the Donaldson matrix. Stokes [32] observed that typically fluorescent emissions arise only at wavelengths that are longer (lower energy) than the excitation wavelength; consequently, the EEM is triangular. It is impossible for us to specify absolute levels for the entries of the EEM; in the simulations, we normalize the EEM so that the maximum value is one. Hence, fluorophore concentrations are estimated in relative units.

#### 3.2.1. Fluorophore mixture model

We modeled five fluorophores that are commonly found in human oral cavity mucosal cells: NADH, FAD, elastin, collagen and porphyrins. The emissions from a mixture is the weighted *o:* _*j*_ sum of these individual terms [33]:

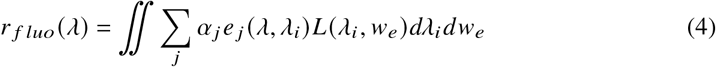

The final expression for the radiant intensity in the output direction, as a function of the fluorophores and standard reflectance is:

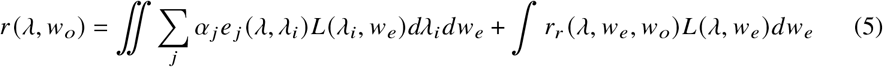

The excitation-emission matrices of the five fluorophores are represented in Figure 3. Simple visual examination suggests that it is unlikely we can find a single illuminant to precisely quantify the concentrations of the five fluorophores; although, some ability to separate out the fluorophores should be possible. For example, notice that the porphyrins have a distinctive spectral emission (Figure 3).

**Fig. 3.**
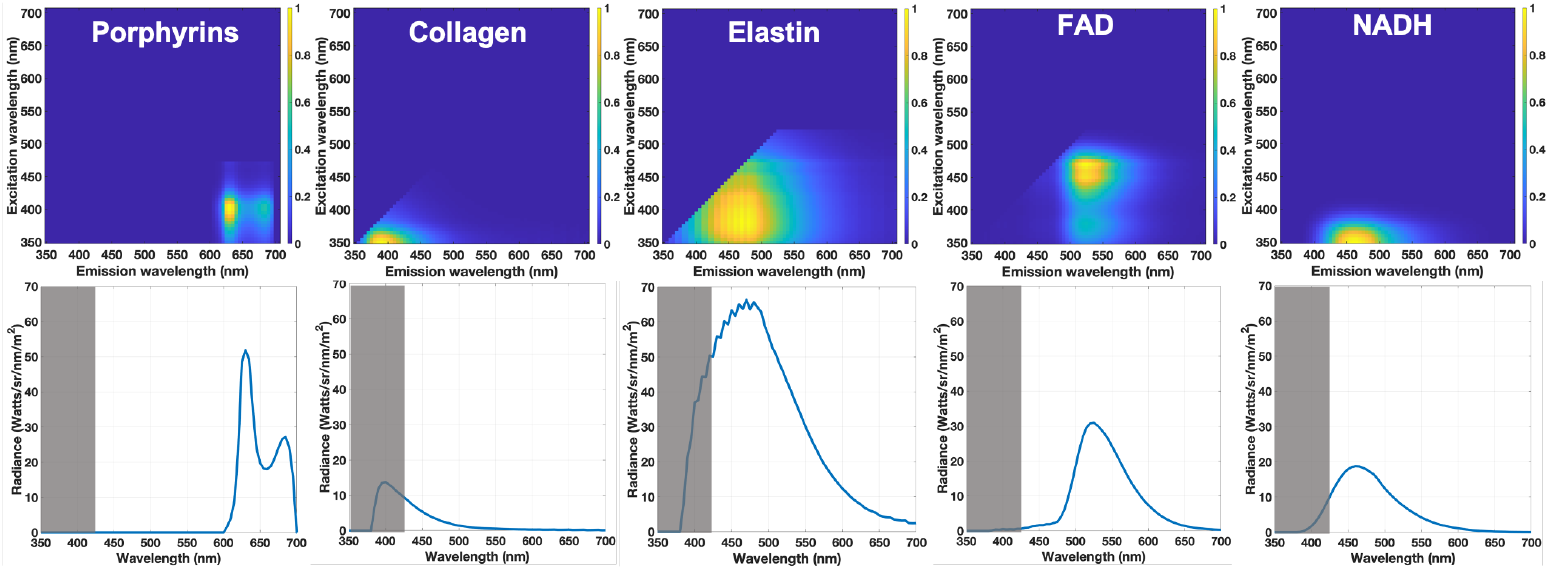
Top row: Excitation and emission matrix for fluorophores commonly found in the oral cavity. The EEMs are triangular because in general the emission wavelength exceeds the excitation wavelength [34]. The emissions are calculated on the assumption that the peak value in the EEM is 1. The gray-shaded region shows wavelengths that are filtered out before reaching the OralEye sensor.

### 3.3. End-to-end simulation of the image system components

#### 3.3.1. Camera lights

The experimental camera (built by FengYun Vision Technologies and referred to as the “OralEye”) is shown in Figure 4. The camera is designed to acquire images for previewing and measuring tissue fluorescence. To meet this goal the camera has two light sources: a ring of LEDs that provides broadband illumination (“white”), and a second array of short-wavelength sources (“blue”) that are used to evoke the tissue fluorescence. The camera acquires images in rapid sequence using different sources with a programmable range of sensor gains and exposure durations. Figure 4 shows the spatial configuration of the two sets of LEDs. In addition, the Figure shows measurements of the relative spectral energy distribution of the white and blue sources.

**Fig. 4.**
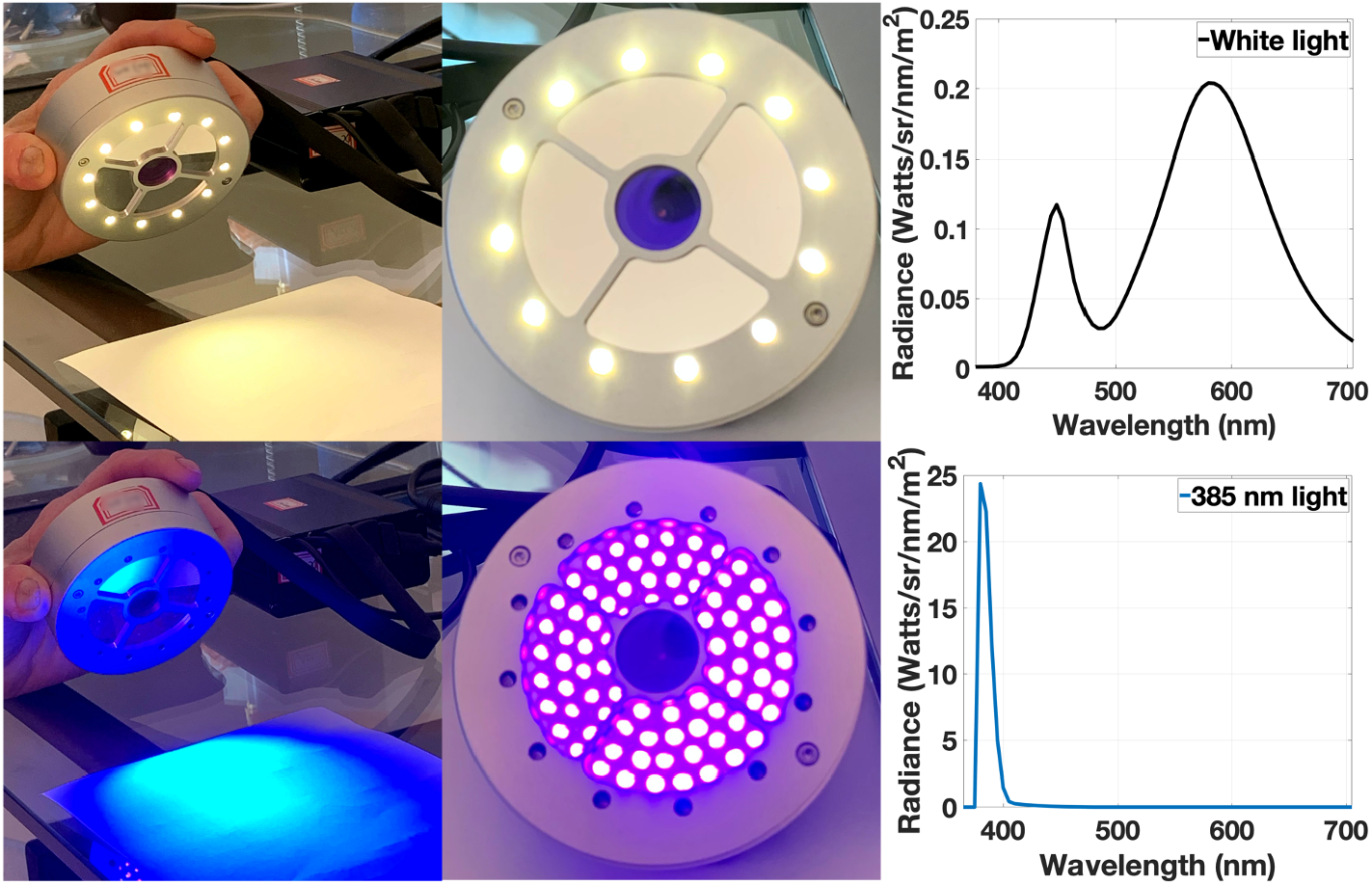
Photos of the OralEye camera, emphasizing the two camera light sources: broadband white light (top row) and blue light (bottom row). The spectral energy of the white is broadband; the blue LED peaks at 385 nm. The white light is used for previewing the oral cavity and the blue light is used for exciting fluorescence. The sensor is in the center of the camera behind a longpass and NIR filter (see Figure 5(c)).

In any realistic setting, it is impossible to precisely control the spatial distribution of the illumination. The illumination will be nonuniformly distributed over space and this nonuniformity will vary with the distance from the camera to the oral cavity. Further nonuniformities arise from secondary bounces of the light from the materials. This spatial illumination nonuniformity is modeled in the simulations and must be accounted for in algorithms that aim to quantify tissue fluorescence from the camera image.

### 3.3.2. Lens and filter selection

An essential aspect of the image system design is to select lights and filters that enable accurate detection of the fluorescent signal from the background of unwanted (diffuse and gloss) light. The fluorescence signal intensity is more than three orders of magnitude lower than the signal level from typical diffuse and gloss reflectance in the oral cavity. The selection of the filters and lights is a significant factor in this design. Even when calibrating the fluorescence signal it was necessary to control the LED spectral distribution to comply with the dynamic range of the PR 670 spectral radiometer.

In order to separate the fluorescence and reflectance signals, the image system includes dichroic filters that limit the wavelengths (a) entering the scene from the blue LED, and (b) entering the camera from the scene (see Figure 5). Specifically, the blue LED sources, with a wavelength centered at 385 nm, emit spectral energy in the wavelength range up to 440 nm that is 2.5 orders of magnitude lower than the peak energy at 385 nm. Even this light level, when scattered by the tissue in the oral cavity, will reduce sensitivity to fluorescence. We placed a dichroic filter (Hoya Y44) in front of the blue illuminants to further reduce the intensity of illuminant energy above 475 nm.

**Fig. 5.**
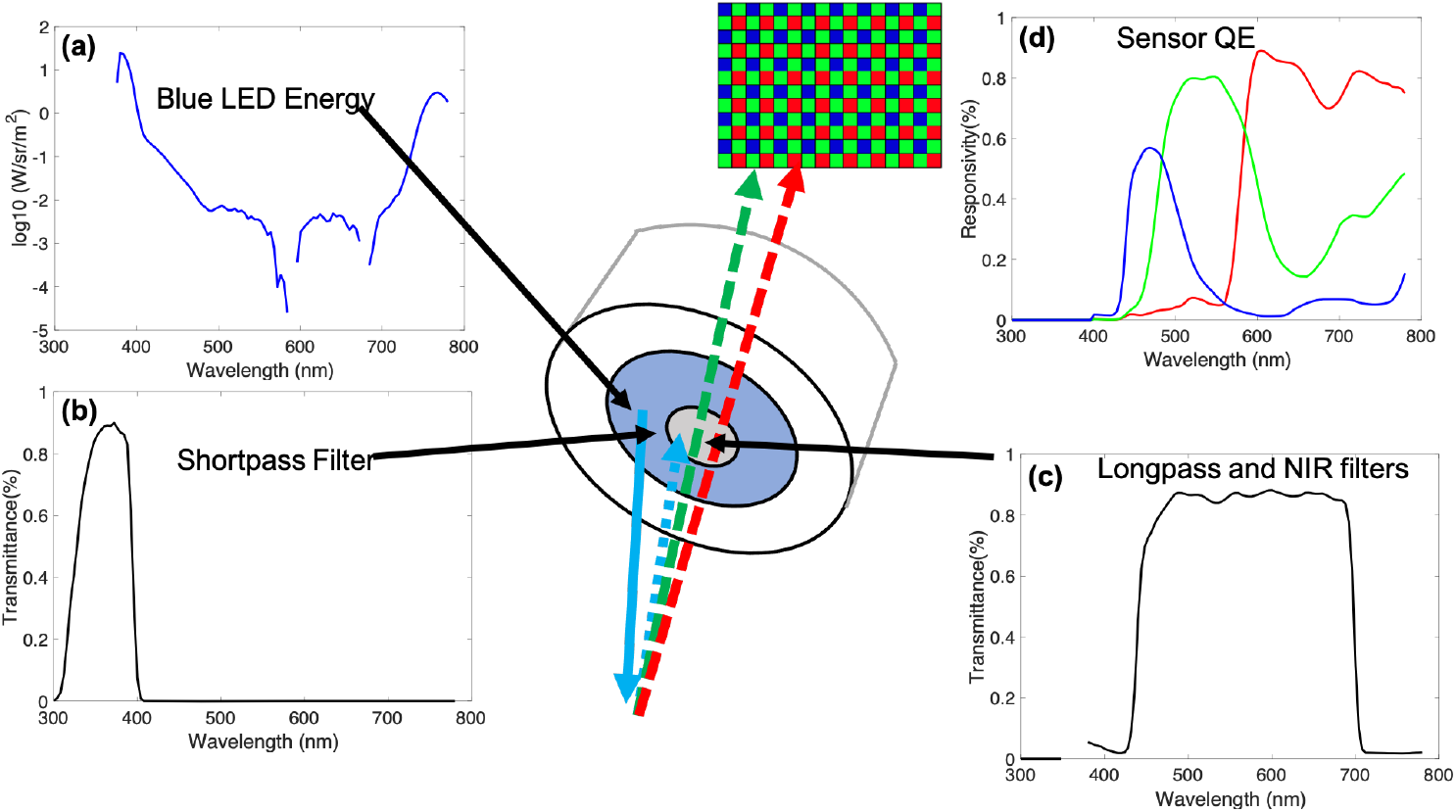
Spectral characterization of the OralEye image system. a) The blue LED spectral energy (plotted on a logarithmic axis) peaks at 385 nm but only drops by 3 orders of magnitude at 450 nm. b) The LED emission band is further narrowed by a shortpass filter which blocks wavelengths > 425 nm. c) A filter in front of the sensor further limits light below 425 nm and longer than 700 nm. This filter is nearly transparent to wavelengths in the range between 450-700 nm. d) Effective sensor spectral responsivity of the three color channels.

The blue LED sources also emit light in the NIR. We placed a second filter in front of the lens to block wavelengths less than 475 nm and greater than 700 nm from reaching the imaging sensor. Hence, under the blue LED illumination conditions, the sensor responds only to irradiance in a range from 475-700nm.

#### 3.3.3. Sensor

The geometric and electrical sensor parameters, including pixel size, resolution (number of pixels), and noise properties are listed in Table 1. We implemented a sensor simulation using ISETCam^2^.

**Table 1.** Sensor geometric and electronic properties.

| Properties | Parameters | Values (units) |
| --- | --- | --- |
| Geometric | Pixel Size | [2.5, 2.5] ($\mu\text{m}$) |
|  | Fill Factor | 0.99 |
| Electronics | Well Capacity | 9000 $e^-$ |
|  | Voltage Swing | 0.911 (V) |
| | Conversion Gain | 0.10 (mV/$e^-$) |
|  | Analog Gain | 1 |
|  | Analog Offset | 0 (mV) |
|  | Quantization Method | 12 (bit) |
| Noise Sources | DSNU | 0.2536 (mV) |
|  | PRNU | 2.29 (%) |
|  | Dark Voltage | 0.15 (mV/sec) |
|  | Read Noise | 1.39 (mV) |

The sensor pixel response can be expressed as:

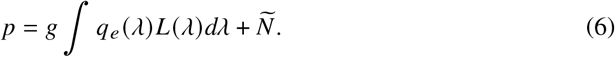

where *p* is the pixel response (*e*^−^), *L* is the irradiance(*q*/*s*/*m*^2^/*nm*), *q*_*e*_ is the sensor spectral quantum efficiency(*e*^−^ /*q*); *Ñ* is noise (*e*^−^); *g* is a scale factor that combines pixel area and exposure time. The values of *g* and *Ñ* are calculated from the irradiance level (quantal noise) and the sensor electrical and geometric properties in the ISETCam simulation. (*q* is quanta, *e*^−^ is electrons, *s* is seconds).

The ISETCam software has been validated in a number of independent experiments [35–37]. For the OralEye device, we validated the sensor model by capturing images of a calibrated color target that included painted surfaces that emit fluorescence when illuminated with the short-wavelength ‘blue’ light [38]. The sensor delivers the raw (linear) RGB data in the Bayer mosaic, which is proportional to the number of electrons. We used bilinear interpolation to demosaic the real and simulated sensor images shown in this paper.

### 3.4. Subjects

Ten participants (seven males, three females; median age 24 years, range 20–67 years) participated in the study, which was approved by the Institutional Review Board at Stanford University. All subjects gave informed consent. The OralEye measurements for each subject took less than one minute, and the exposure to the blue LED was approximately 30 ms. Spectrophotometric measurements of tongue radiance were obtained from two of the participants. These measurements, including setup time, took approximately ten minutes and the exposure to the blue LED was approximately 10 sec. All of the exposures were well within the safety limits for exposure to short wavelength light.

## 4. Results

Renderings from the end-to-end simulation of the oral cavity through the OralEye camera are compared with real OralEye images in Figure 6(a-f). The images compare the measured and expected results when using the blue LED. The simulation models fluorescence of both the oral cavity and teeth [39–42].

**Fig. 6.**
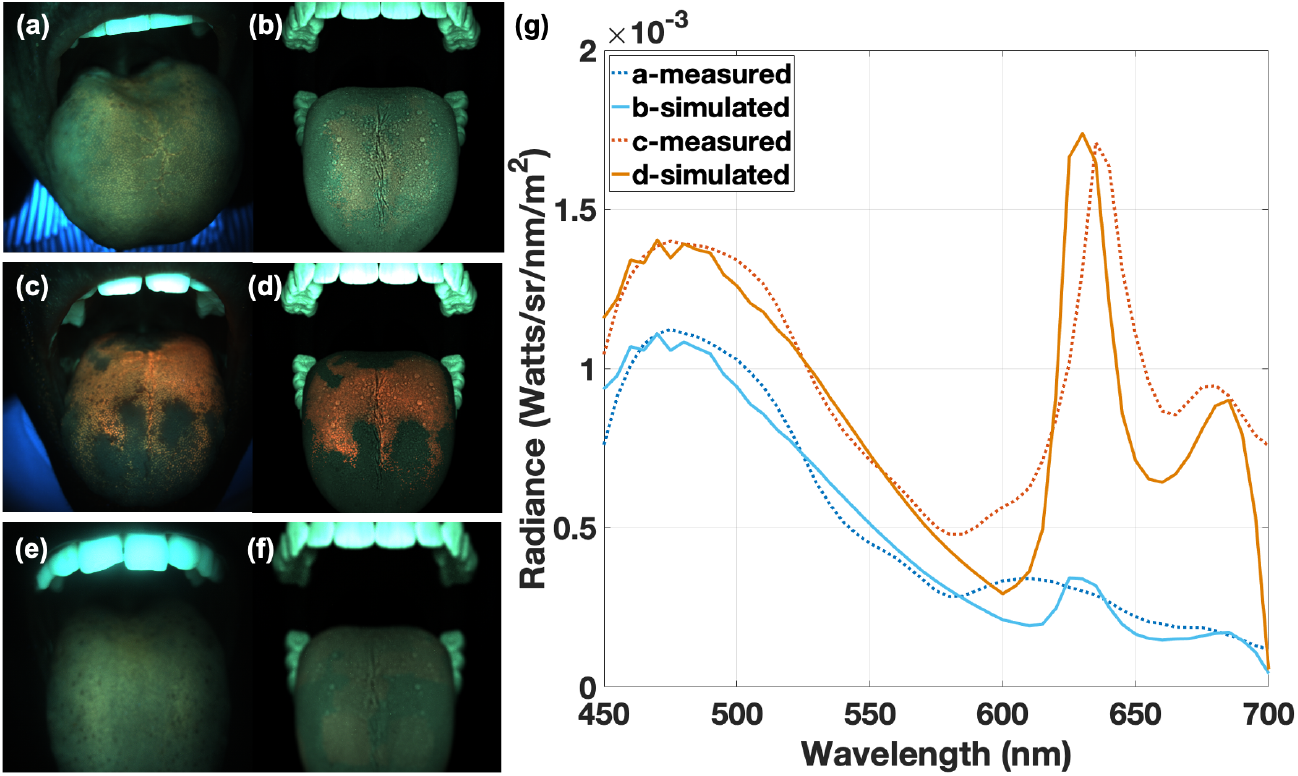
Comparison between end-to-end simulations and OralEye images of the oral cavity in three participants. The images reflect the typical variation in the porphyrin concentration among healthy participants. Panels (a, c, e) show OralEye images and panels (b, d, f) show the corresponding simulations. (g) Comparison between measured (dotted) and simulated (solid) radiance of two points on the dorsal tongue from high and low porphyrin regions in two participants.

Tongue fluorescence was modeled by a weighted combination of NADH, FAD, collagen, elastin and porphyrin emissions. Figure 6(g) compares the spectral radiance of simulated and measured fluorescence from two points on the dorsal surface of the tongues in two participants. The measurements were obtained using a PR670 spectroradiometer equiped with a longpass filter (Hoya Y44) that blocked light below 425 nm. Without this filter, the fluorescence would have been too weak to detect given the dynamic range of the spectroradiometer. The simulated radiance in Figure 6(g) was also filtered by the same longpass filter.

We refer to the combined signal from NADH, FAD, collagen, and elastin as the “bulk fluorescence”. There are many possible combinations of these four fluorophores that predict the measured radiance. For this reason, using the 385 nm LED alone does not enable separating (spectrally unmixing) the fluorophores in the bulk fluorescence signal. As a summary of the multiple solutions, we can say that the combinations that predicted bulk fluorescence measurements in two participants (see Figure 6(g)) had zero concentration of NADH, low levels of FAD and collagen, and high levels of elastin.

The porphyrin concentrations were chosen to approximate the measured spectral radiance, which differed for the two participants. The region measured for the participant in panel (a-b) was predicted mainly from the bulk fluorescence with a small amount of porphyrins. The region measured for the participant in panel (c-d) was predicted from the bulk fluorescence with a high concentration of porphyrins.

To generate realistic spatial distributions of porphyrins, we utilized the additivity property of sensor responses from different fluorophores. We simulated two sensor images: one simulates the bulk fluorescence with no porphyrin, the second simulates porphyrin emissions with no bulk fluorescence. For each participant we created a spatial mask that indicates the locations with a significant porphyrin concentration. The final rendered image is the sum of the porphyrin sensor image multiplied by the mask and the bulk fluorescence image.

The specific geometry (overall size and shape, distinctiveness of the teeth, tongue position) differ, but the general color in the images and the properties of the nonuniform illumination are similar. A property shared by the simulations and the real images is that the absolute level of the digital values depends significantly on spatial variations in the illumination. This suggests that fluorophore estimates should be based on the relative, not absolute, RGB values.

### 4.1. Sensor chromaticity

The significant illumination variation in the scene contributes to the varying fluorescent emissions at different locations, eliminating the opportunity to use absolute RGB levels to measure fluorescence. Because we rely on the ratio of the RGB values, the chromatic information about the fluorophores is two-dimensional. Furthermore, the literature informs us that there are a large number of different fluorophores that can appear in the mucosal tissue in different combinations (Figure 3). The simulations show that different combinations of tissue fluorophores may produce the same spectral radiance and, consequently, the same sensor responses.

For this system, there is one possible source of meaningful chromatic information. The porphyrins EEM differs substantially from the other principal fluorophores. Consequently, a strong porphyrins signal has an impact on the RGB values that can be distinguished from the other fluorophores.

We simulate the expected impact of porphyrins, as measured in sensor chromaticity space, in Figure 7. Figure 7(a) illustrates the simulated R and G values assuming two different groups of fluorophores. The green points represent a noisy signal from a bulk mixture of the FAD, collagen and elastin. The red points represent the signal expected from porphyrins, again assuming a particular concentration and illumination level. In both cases, the measurements might fall anywhere along the two dashed arrows depending on the fluorophore concentration and local illumination level. In this simulation we plot the data as if there is no diffuse or glossy light reflected from the tissue, though in reality both of the lines would start from a small non-zero position in the graph in the presence of the weak blue light reflected from the tissue. The contribution of the reflected light is relatively small because the filters remove most of the diffusely reflected light which is below 425 nm. In the complete simulation and when comparing with the measurements, we account for the light reflected from the tongue.

**Fig. 7.**
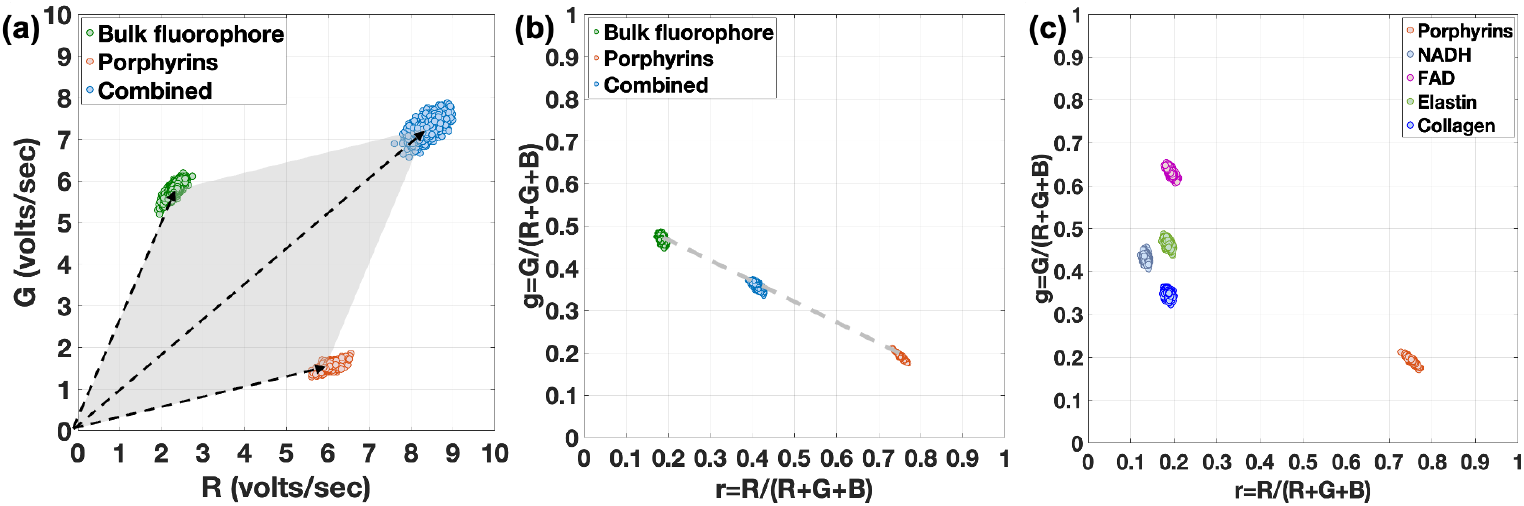
Illustration of fluorescence sensor responses and sensor chromaticities. a) R and G sensor response representation. The points plotted in green and red are expected R,G signals, including noise, from the bulk fluorophore and porphyrins respectively. The signal expected from the mixture of these fluorophores is the vector sum of these two points, plotted in blue. For different relative amounts of the two fluorophores, the mixture will fall within the grey-shaded parallelogram. b) Sensor chromaticity representation. The sensor responses are normalized across three channels to eliminate the impact of non-uniform illumination and fluorophore concentration. The sensor chromaticities of the combined fluorophores will fall along the line connecting the chromaticities of the bulk fluorophore and porphyrins. The position along the line will depend on the relative strength of the two signals. c) Sensor chromaticities of the individual fluorophores. Each fluorophore is represented at a different location on the sensor chromaticity graph. For the excitation light of 385 nm and the OralEye spectral sensitivity, the porphyrins are widely separated from the cluster of the other four fluorophores.

Depending on the fluorophore concentrations and illumination level, the combined signal might fall anywhere in the gray shaded region. For example, when the fluorophores are present at the concentrations indicated by the red and green line endpoints, the combined signal will be located at the tip of the blue shaded region. The position within the gray shaded region provides information about the relative amount of the bulk and porphyrin fluorophores.

Figure 7(b) represents the same information but plotted with respect to sensor chromaticity values (r,g). The sensor chromaticities are the R (G) values divided by the sum of the R, G and B values:

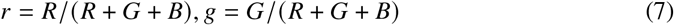

From the formula, we can see that all RGB-values that fall along a line *α R, G, B* share the same sensor chromaticity. The reason for representing the data with respect to sensor chromaticity is that the value is invariant with respect to the absolute fluorophore concentration and absolute illuminant intensity, two factors that we cannot control. Mixtures of the two signals *r*_1_, *y*_1_ and *r*_2_, *y*_2_ will fall along a line between the two chromaticities (dashed line, Figure 7(b)). The position on the line will depend on the relative intensity of the two emissions.

The expected positions in sensor chromaticity space of five different oral cavity fluorophores are shown in Figure 7(c). For this camera the sensor chromaticities of most of the fluorophores fall in a small region of the sensor chromaticity plane. The proximity of these values, coupled with the metamerism described earlier, makes it difficult to discriminate the relative contributions of these fluorophores. The porphyrins contribution, however, is relatively distance and has the possibility of drawing the total signal away from the cluster. This is a feature of the simulation that we can confirm with respect to the measured images.

### 4.2. Validation with empirical measurements

Figure 8(a-c) shows the OralEye images captured from the dorsal tongues of three healthy individuals as shown in Figure 6. These images were captured in a dark room using only the blue LED illuminant. Because of the filters, the captured light is almost entirely fluorescence; the reflected light is mainly confined to short wavelengths below the acceptance region of the camera. The teeth are very fluorescent and emit light over a wide range of wavelengths.

**Fig. 8.**
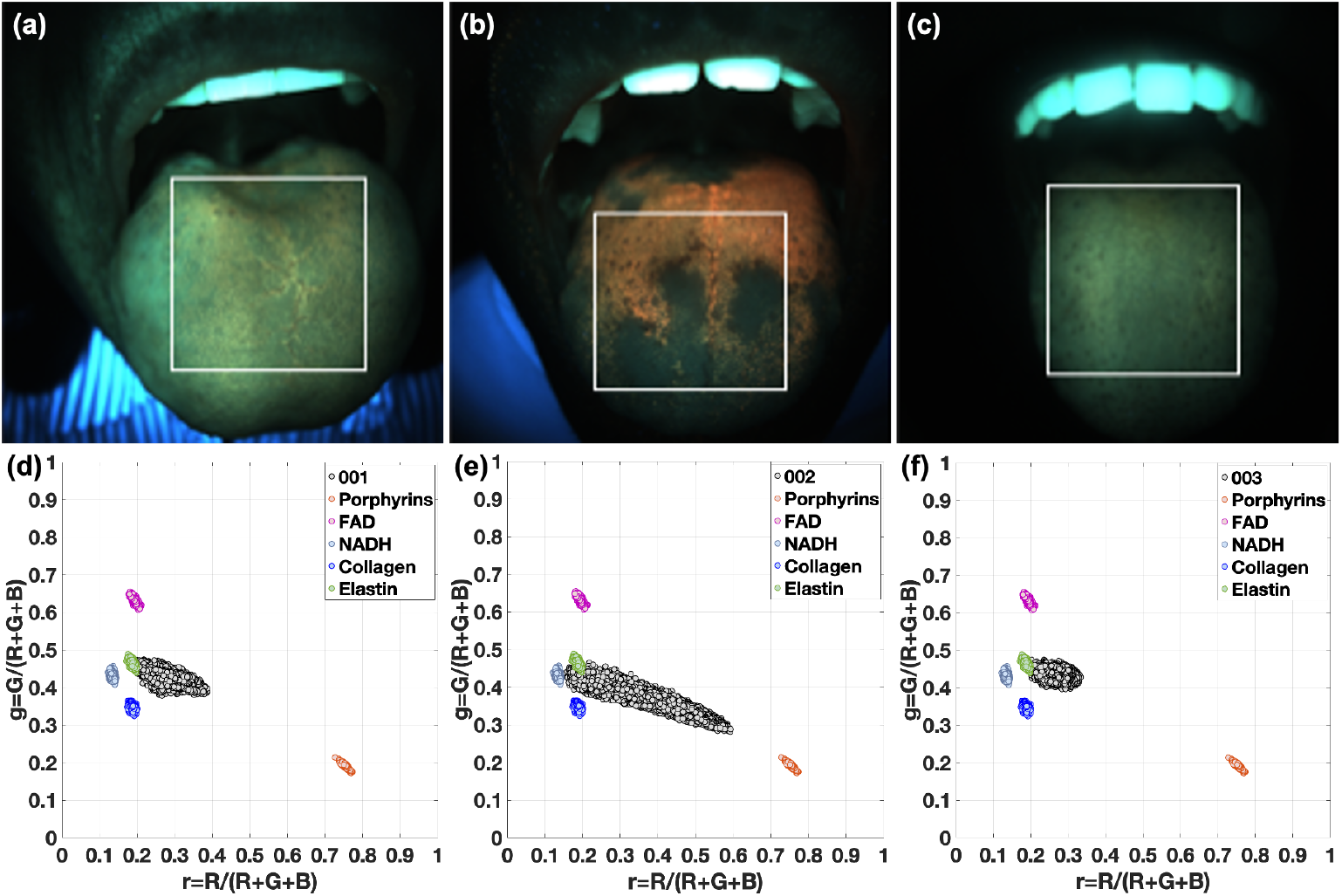
Sensor chromaticity analysis of OralEye images of three participants. The sensor chromaticity values of pixels in the regions of interest denoted by the white rectangles are plotted as black points in the bottom row. The data fall along a line extending from the position of elastin extending to the position of the porphyrins. The length of the line and its endpoints differ between the subjects. The sensor chromaticity data in the middle column extend further to the left than the data in the left and right columns.

We analyzed the sensor chromaticity in the OralEye images and compared them with the values expected from the simulation. Specifically, we selected a large region within the dorsal tongue (white box), and we plot the sensor chromaticity coordinates for all the pixels in this region (Figure 8(d-f)). The sensor chromaticity values are shown along with the simulated values for the different fluorophores (Figure 7(c)). The sensor chromaticity data align well with the expected chromaticity values, falling along a line that extends from the central position of the bulk fluorophores (NADH, FAD, collagen and elastin) in the direction of the porphyrin fluorophore. The fact that the sensor chromaticity values follow a similar pattern for all participants confirms (a) the accuracy of the simulations, and (b) the expectation that the primary difference we observe is explained by the porphyrins concentration.

Porphyrins fluorescence is measured on the dorsal side of healthy tongues illuminated with short wavelength light [20]. We observed no porphyrins emissions from the sides of the tongue, the ventral surface, or on the upper and lower palates. Differences in the amount of porphyrin signals on the dorsal tongue of different participants may be due to diet and the time of day. For example, the OralEye camera image shown in Figure 8(b) on was captured from the tongue of an individual who had recently eaten lunch. The OralEye image in Figure 8(c) was captured from the tongue of a vegetarian just before lunch. In a separate measurement, we observed porphyrins fluorescence on the tongue of this same individual just after drinking mango juice.

We collected data from ten subjects, and the general agreement in the sensor chromaticity properties from this admittedly small number of participants is encouraging. It suggests that we might be able to define a narrow, quantitative expectation for the chromaticity range in the healthy dorsal tongue. On the other hand, the data also suggest that with this camera design, it will be impossible to estimate the relative proportions of elastin, FAD, collagen, and NADH fluorescence from measurements of the bulk fluorescence.

## 5. Discussion

For the last twenty years, digital camera design has been driven by consumer photography applications. Hardware and software components have been optimized to capture radiance signals that humans can perceive, and the camera image processing pipeline is designed to produce images that appear to be pleasing to consumers [43–45].

The first implementation of the OralEye image system uses hardware components that were developed for consumer photography; but the system has a different purpose. The system is intended to quantify the amount and type of tissue fluorescence in a large field of view within the oral cavity that is invisible to humans under normal viewing conditions. Consequently, the system design integrates special purpose illuminants, filters, and sensors that are outside of the usual scope of consumer photography. The images the system produces are not intended for consumers or clinicians to view, but rather for clinical laboratory tests that quantify the fluorophore concentrations in the oral cavity.

The simulations show that the current system design can estimate the combined emissions from relatively high concentrations of certain fluorophores (collagen, elastin and FAD), which we refer to as “bulk fluorescence”. These fluorophores produce higher G sensor values than combinations that have lower concentrations of these tissue fluorophores. The system has only one excitation light, and the simulations reveal that several different concentrations of these fluorophores produce the same bulk fluorescence. Hence, it is not possible to determine the relative concentrations of these tissue fluorophores by analyzing the sensor data from the current system. The porphyrins, however, stand out because their fluorescence signal dominates the R sensor values. Hence, the system can measure the relative balance between the bulk fluorescence and the porphyrins. We confirm this ability using both simulations and experimental measurements.

The image systems simulations have helped us both understand and quantify the interaction between fluorophores, illuminant spectra and measured fluorescence. The validation of the simulations encourages us to use the software to explore new image system designs.

### 5.1. Design considerations

The first challenge we confronted in this design was to eliminate the impact of the reflected light. The reflected light was as much as four orders of magnitude more intense than the fluorescence emitted by oral mucosal tissue. As the measurements show, the system is inadequate to simultaneously measure the reflected and fluorescent components. We excluded the reflected light by placing a shortpass filter in front of the 385 nm LEDs, blocking light energy in the longer wavelengths from reaching the oral cavity. Selecting the light and filter was an essential part of designing the system.

A second challenge arises from the inability to illuminate the oral cavity uniformly. The complexity of the illuminant shading is due in part to the geometry of the lights, but it is also due to the fact that the oral cavity is a three dimensional structure with surfaces at different depths that can occlude and cast shadows on other surfaces. The sensor RGB values from nearby regions on the same surface may differ because the light is non-uniformly distributed over the surface, the orientation of the surface, or the amount of indirect lighting. We suspect that this issue will persist through all system designs, and for this reason an approach based on sensor chromaticity may continue to prove helpful.

A third challenge we will confront is how to separate the signals within the bulk fluorescence. Through the validated simulation methods, we are exploring designs that include multiple excitation wavelengths and commercial multispectral sensors.

### 5.2. Applications

The ability to quantify the relative amounts of porphyrin and bulk fluorescence may benefit several applications in dentistry. For example, porphyrins fluorescence is generated by bacteria that accumulates on teeth and dentures [27], in crevices [28], and along the gum lines [46, 47]. This observation led to the development of adjunct dental devices that use fluorescence imaging to help dentists visualize the location of bacteria associated with caries and gingivitis [48, 49]. For these applications, there may be value in using an imaging system that can document the location and quantify the relative concentrations of porphyrin in different parts of the oral cavity.

The porphyrins fluorescence from the dorsal surface of tongues in healthy individuals [20] is attributed to a complex community of bacteria referred to as the oral microbiome [50]. Analysis of the tongue dorsum microbiome is understudied, particularly when compared to the amount of research devoted to studying the gut microbiome [51]. Monitoring and manipulating the oral microbiome will lead to a better understanding of the functional role that oral bacteria have on the dorsal surface of the tongue.

Dentists also use adjunct devices to visualize bulk fluorescence in oral mucosal tissue. Clinicians are trained to look for dark areas where bulk fluorescence is not visible as an indicator of the degradation of the structural integrity of tissue or changes in tissue metabolism. The assessment is visual and subjective and thus the efficacy of these devices depend on clinician experience. These devices may help clinicians find areas they might otherwise overlook, but they do not help them differentiate between dysplasia and benign inflammatory conditions [15]. Consequently, the sensitivity of these devices is high, but the specificity is low [52, 53].

A main goal of our work is to design an image system that replaces the subjective judgments of a clinician with a lab test that meaningfully assesses the health status of the oral cavity. We have shown that the OralEye camera can quantify the relative amounts of porphyrin and bulk fluorescence in healthy subjects. Assuming that NADH emissions are negligible, decreases in bulk fluorescence may indicate degradation of the structural integrity of tissue (associated with decreases in elastin and collagen) or changes in tissue metabolism (associated with a decrease in FAD). To determine whether these measurements are diagnostic, it will be necessary to collect additional data from patients that have dysplasia and cancerous lesions.

### 5.3. Future Work

We are extending the work we describe in this paper in two ways. First, we are using the current OralEye camera to collect additional data in both healthy individuals and patient populations. To pursue these measurements, we automated the data storage and analysis using a cloud-based data management system (Flywheel.io). This system anonymizes the data while at the same time storing important demographic information. The normative data that we are collecting in healthy individuals will define a distribution against which we can compare the data captured from the patient population. Aggregating these data and monitoring patient outcomes, should enable us to improve oral health predictions. Ultimately through the acquisition of quantitative data about the fluorescent signals from clinical cases, we may be able to implement meaningful diagnostic tools.

Second, we are using image systems simulation software to create soft-prototypes of multispectral imaging systems that combine multiple illuminants with multiple imaging sensors. We will use sensor chromaticity values to quantify the relative concentrations of different tissue fluorophores. The simulations will enable us to determine whether it is possible to design multispectral imaging systems that can provide information about the relative concentrations of NADH, FAD, collagen and elastin and to predict the efficacy of the soft-prototypes before building a real physical device.

## 6. Conclusion

Image systems simulations enable us to create software prototypes of digital cameras and to predict the data we would capture for different combinations of tissue fluorophores, illumination and imaging sensors. We describe and provide open-source freely available software prototyping tools that can be used to design and evaluate new imaging systems based on multiple lights and novel imaging sensors^3^.

We used image systems simulations to design an imaging system capable of exciting and measuring fluorescence in the oral cavity. We created a hardware prototype of the imaging system (OralEye) and compared the data we collected from the real device with the data we predicted from the software prototype. The simulations and data suggest that sensor chromaticity values, derived from real and simulated OralEye RGB camera data, are useful for estimating quantities that are invariant to changes in the spatial distribution of lighting. Specifically, the sensor chromaticity values can quantify the fluorescence due to porphyrins relative to the combined emissions from other fluorophores in the oral cavity, referred to as the bulk fluorescence. Additional data from patient populations and from different regions of the oral cavity should prove informative as to the diagnostic value of the porphyrin and bulk fluorescence estimates.

## Acknowledgements

We thank Rangtao Huang, Tanglong Wang, and Xixi Li at FengYun Vision Technologies for software and hardware support of the experimental camera (OralEye). We thank Henryk Blasinski, Zhenyi Liu, Kaijun Feng and Krithin Kripakaren for their contributions to the simulation software and camera assembly. We thank Adam Wandell, Chris Holsinger, Tulio Valdez and Thomas Goossens for many helpful discussions and feedback about this project.

## Footnotes

1 https://github.com/ISET/iset3d/wiki

2 https://github.com/ISET/isetcam/wiki

3 https://github.com/ISET

